# Bioinformatic analysis reveals mechanisms underlying parasitoid venom evolution and function

**DOI:** 10.1101/423343

**Authors:** Gloria Alvarado, Sarah R. Holland, Jordan DePerez-Rasmussen, Brice A. Jarvis, Tyler Telander, Nicole Wagner, Ashley L. Waring, Anissa Anast, Bria Davis, Adam Frank, Katelyn Genenbacher, Josh Larson, Corey Mathis, A. Elizabeth Oates, Nicholas A. Rhoades, Liz Scott, Jamie Young, Nathan T. Mortimer

## Introduction

Parasitoid wasps are a numerous and diverse group of insects that obligately infect other arthropod species. These wasps lay their eggs either on the surface or within the body cavity of their hosts, and the resulting offspring exploit the hosts’ resources to complete their development [1]. Many parasitoids also introduce venom gland derived proteins or polydnaviruses into the host during infection. These factors act through a variety of mechanisms to manipulate host biology in order to increase the fitness of the developing parasitoid offspring [2–4]. Many hosts mount immune responses to parasitoid infection, and accordingly, parasitoids have evolved multiple venom genes that encode immunomodulatory factors. These factors target host cellular and humoral immune mechanisms, to allow the developing parasitoid to evade the killing activity of host immunity [5–11]. These parasitoid immunomodulatory activities are strikingly diverse, and include venoms that block host immune cell signaling [5,6], alter host gene expression [11,12], or even cause immune cell death [13]. Wasp venoms also contain proteins that manipulate host physiology and metabolism [14–16], including factors that alter host development [17,18], lipid metabolism [19], and nutrient flux [20–22]. These manipulations are thought to improve the nutritional content of the hosts’ internal environment in order to benefit the developing wasp [2]. Along with these physiological targets, many parasitoids can also use venom factors to alter their hosts’ behavior [23–28]. These modified behaviors promote parasitoid survival and development; often at the expense of the hosts’ own fitness.

Most molecular studies aimed at uncovering the functions of parasitoid venom proteins have been focused on the characterization of a single protein. However, with the advent of high throughput “venomics”, venoms can now be considered as a whole in order to gain insight into higher order venom properties. These venomic studies have used an approach that combines genomic or transcriptomic sequencing with the purification and mass spectrometry based identification of venom proteins. This approach allows for the entire venom protein content to be identified in a single experiment, and has been completed on diverse parasitoid wasp species [5,29–38]. As the number of venomics projects has increased, understanding the composition of venoms has become an active area of research. This includes studies focused on exploring both the evolution of venom content and venom protein function [15,39].

The evolution of venom content is an especially interesting question in evolutionary biology. It has been hypothesized that proteins may be recruited into the venom proteome through a variety of processes, including: gene multifunctionality, in which genes are active in parasitoids in both cellular and venom contexts [39]; the co-option of cellular genes into venom specific expression and function [40]; the evolution of alternative splicing products of cellular genes that result in venom specific protein isoforms [5,41]; gene duplication events that create venom specific paralogs [29–31]; the evolution of *de novo* genes [39]; and the horizontal transfer of genes between organisms [42]. The ability to study a whole venom proteome will allow us to address venom evolution from a unique perspective, and to test whether these different mechanisms may collectively function to drive the evolution of venom composition within a single parasitoid species. Similarly, while the “gene-by-gene” approach has revealed the molecular details of venom protein function, the consideration of predicted functions of the venome as a whole can provide additional insight into understanding venom function.

Due to the wide range of encoded activities, understanding the composition and function of parasitoid venoms is also a subject of great interest in applied biology. Parasitoids are important biological control agents for integrated pest management programs [18,43], and as a result of their ability to disrupt the development of insect pests, parasitoid venom factors may be useful as the basis for the synthesis of novel bio-insecticides [2,15,18]. Furthermore, many parasitoid venoms specifically target conserved signaling pathways in their hosts, including the Toll and JAK-STAT signaling pathways, intracellular calcium signaling, and Rho family GTPases [5,6,12]. Since homologues of these pathways play conserved roles in human health, these venom proteins may have the potential to be developed as pharmaceutical agents [2,15,44].

We are particularly interested in characterizing the venoms of the *Drosophila*-infecting parasitoid wasps. These wasps are endoparasites that infect *Drosophila* species during the larval stage, and complete their own development during the fly larval and pupal stages [1]. Since they develop within the larval body cavity, these parasitoid species encounter the fly’s robust cellular immune response, and accordingly have evolved virulence proteins that allow them to evade fly immunity [5–7,45–48]. We have previously identified 166 venom protein encoding genes from the *Drosophila*-infecting parasitoid *Ganaspis sp.1* [5]. Here we take a whole venom bioinformatic approach to analyze the evolution as well as putative functions of *Ganaspis sp.1* venom.

## Methods

### Sequence Data

The *Ganaspis sp.1* sequence data has been deposited under Transcriptome Shotgun Assembly (TSA) project accession GAIW00000000. Other sequence accession numbers were EU482092.1 (*Blastocystis* 18S rRNA), AB038366.1 (*Arsenophonus* 16S rRNA), AB915783.1 (*Sodalis* 16S rRNA), and as given in the text.

### Transcript Abundance

The relative abundances of venom and cellular mRNA transcripts in our RNAseq data were compared using the RSEM software [49]. RNAseq reads were aligned to our *de novo* transcriptome assembly to generate transcript counts in transcripts per million (TPM). TPM values were then ranked and the venom and cellular transcript groups were compared by Wilcoxon signed-rank test (implemented in R).

### Protein Properties

We first generated a random list of cellular proteins using a custom Perl script. The molecular weight (MW) and isoelectric point (pI) of each protein was then calculated using the ExPASy Compute pI/Mw tool [50]. Values were then compared by Welch Two Sample t-test (implemented in R).

### Homology Finding

To find homologous sequences, each *Ganaspis sp.1* venom protein was aligned to the non-redundant database using blastp (accessed February 2016) [51]. Conserved domains were identified using the Batch CD-Search tool to query the Conserved Domain Database (CDD), and the numbers of occurrences of each CDD accession number were counted [52,53]. Domain enrichment was determined using Fisher’s exact test, and domain numbers were compared by Welch Two Sample t-test (implemented in R).

### Phylogeny

To build phylogenies for investigating potential horizontal gene transfers, homologous sequences were identified by homology finding and then processed using MEGA [54]. These sequences were first aligned and the alignments were used to construct Maximum Likelihood phylogenetic trees, with 500 Bootstrap replications and JTT substitution model.

## Results and Discussion

### Characterization of *Ganaspis sp.1* venom encoding genes

We predict that *Ganaspis sp.1* venom protein encoding genes are distinct from cellular (non-venom) protein encoding genes. We expect that when compared to a subset of cellular genes, venom genes would have distinct properties that include expression level and the properties of the encoded venom proteins. We hypothesize that venom encoding transcripts will be among the most abundant, due to the crucial role of the venom in facilitating successful parasitization [40]. To test this hypothesis, we compared the abundance of venom protein encoding transcripts with wasp cellular gene transcripts. We found that venom gene transcripts have significantly higher expression levels, with a mean abundance of 145.15 transcripts per million (TPM), compared to 35.89 TPM for cellular transcripts (Supplementary Table 1; Wilcoxon rank sum test: W = 4020800, p < 2.2e-16).

To test whether the venom proteins from *Ganaspis sp.1* also show distinct physical properties, we compared the predicted molecular weight (MW) and isoelectric point (pI) of *Ganaspis sp.1* venom proteins with a randomly derived list composed of an equal number of *Ganaspis sp.1* cellular proteins. We found that, on average, *Ganaspis sp.1* venom proteins are significantly smaller (MW of 44.3 kDa for venom proteins compared to 62.1 kDa for non-venom proteins, p = 0.004397, Figure 1A), and have a significantly lower pI (6.99 for venom proteins compared to 8.06 for non-venom proteins, p = 1.112e-06, Figure 1B) than *Ganaspis sp.1* cellular proteins. Interestingly, *Ganaspis sp.1* venom lacks the abundant class of small, highly charged peptides that characterize the venoms of many social hymenopteran species [55–59]. These data suggest that the composition of parasitoid wasp venom may be distinct from both cellular proteins and also from the venoms of non-parasitoid hymenopterans.

**Figure 1.**
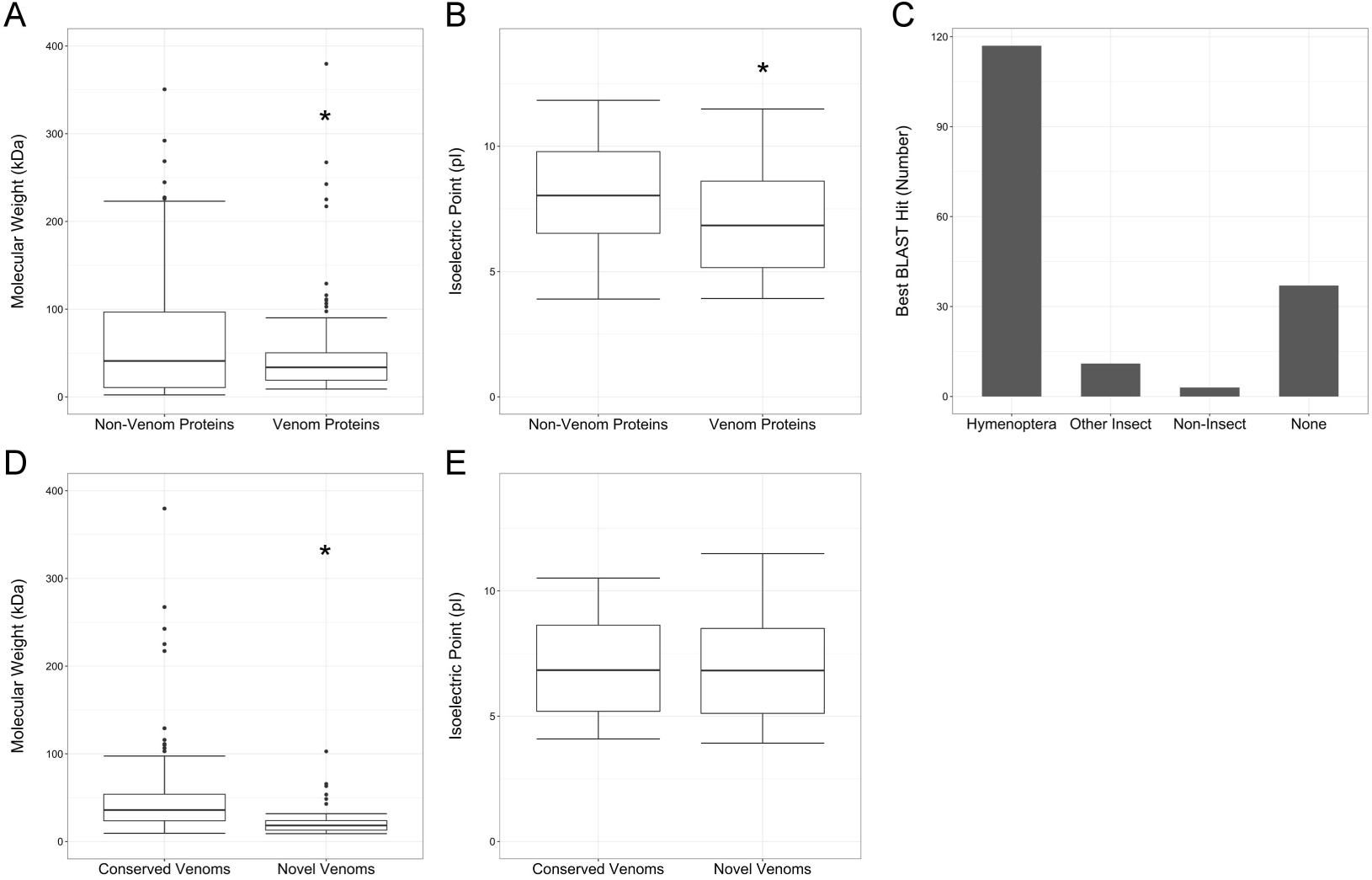
Venom protein characterization. (A,B) The molecular weight (A) and isoelectric point (B) of venom proteins (right) compared to a random subset of non-venom proteins (left). (C) The phylogenetic origin of the the best BLAST hit for each venom protein. *Hymenoptera* indicates that the closest homolog is from an insect within the order Hymenoptera, *Other insect* indicates that the closet homolog is from an insect from any other order, *Non*-*insect* indicates that the closest homolog is from a non-insect species, and *None* indicates that there were no homologous proteins found in the NCBI database. (D,E) The molecular weight (D) and isoelectric point (E) of novel evolutionarily novel venom proteins (right) compared to venom proteins showing evidence of sequence conservation (left). ^⋆^ indicates p < 0.05 by Welch Two Sample t-test.

### Origin of *Ganaspis sp.1* venom encoding genes

Our data can provide insight into the origins of the *Ganaspis sp.1* venom proteome. To investigate the origins of *Ganaspis sp.1* venom proteins, we first used BLAST to identify the closest homolog of each venom. We determined that for 117 venom proteins the closest homolog is found within another hymenopteran species, and that other insect species account for the nearest homolog of a further 11 venom proteins. These findings suggest that the majority of *Ganaspis sp.1* venom proteins are either multifunctional or have evolved from cellular genes (Figure 1C). Interestingly, we also detected 37 venom proteins that do not have homology to any sequence found in the NCBI database (Figure 1C), suggesting that these venom proteins (Supplementary Table 2) may have evolved as *de novo* protein encoding genes. This finding is in accordance with the venoms of other parasitoid species, notably *Nasonia vitripennis*, in which 23 of the 79 identified venom proteins have no sequence similarity to known proteins [2]. The proteins encoded by these putative *Ganaspis sp.1 de novo* venom genes are significantly smaller than the conserved venom proteins (24.8 kDa vs 49.7 kDa, p = 1.345e-05, Figure 1D), but we do not find a difference in pI (6.97 vs 7.00, p = 0.9447, Figure 1E). This smaller protein size is consistent with *de novo* proteins in a wide range of other species, including other insects [60,61], humans [62,63], and plants [64,65].

Our BLAST data further identified 3 venom proteins that only show significant homology to non-insect sequences, suggesting that the genes encoding these proteins may have been introduced into the *Ganaspis sp.1* genome, and subsequently into the venom proteome, by horizontal gene transfer (HGT; [39]). HGT is characterized by the transfer of DNA between species, and HGT events can be seen as phylogenetic incongruities [66,67]. The first of these non-insect derived genes is *comp112* which shows homology to the *AV274*_*2271* protein from *Blastocystis sp. NandII* (Figure 2A). *Blastocystis spp.* are single celled intracellular protozoan parasites that infect a wide range of species from humans to insects [68,69]. The other two non-insect derived venom genes, *comp3581* and *comp11645*, are most similar to genes from insect symbiotic enterobacteria species [70,71]. *comp3581* is homologous to the *WP 032114503.1* gene from *Arsenophonus sp.* (an endosymbiont of *Nilaparvata lugens*; Figure 2B) and the *comp11645* gene is homologous to the *SopA*-*like* gene from *Sodalis sp.* (Figure 2C).

**Figure 2.**
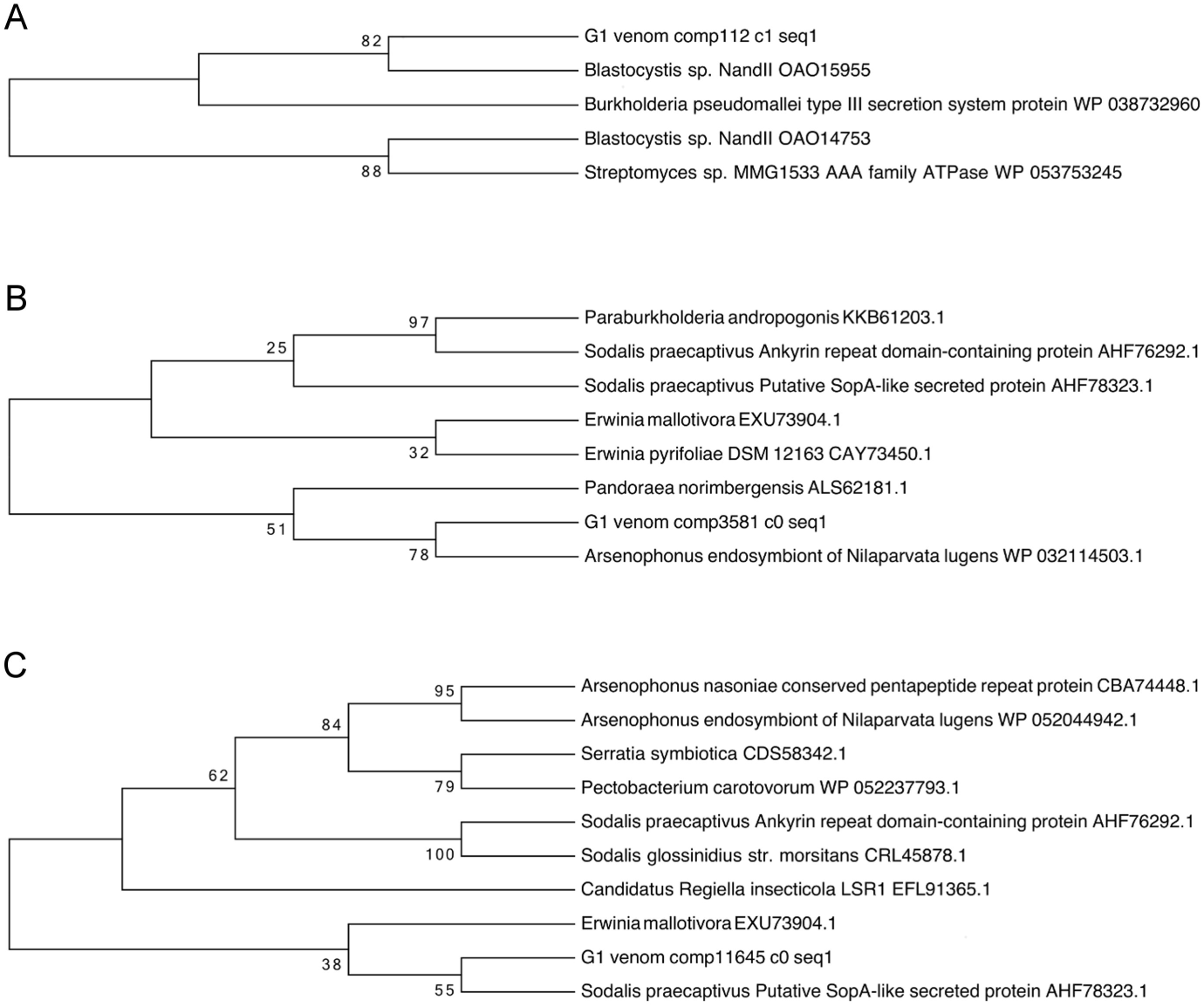
Phylogenetic analysis of putative horizontally transferred genes. (A-C) Phylogenetic trees showing the closest homologs of the venom proteins from the *Non-insect* class of BLAST results: (A) *G1 venom comp112*, (B) *G1 venom comp3581*, (C) *G1 venom comp11645*. Numbers represent bootstrap values (with 1000 replicates).

It has recently been noted that many proposed examples of HGT events may instead represent sequencing artefacts [72,73]. However, due to our workflow, we have isolated these 3 putative horizontally transferred genes from distinct biological samples; the transcripts were sequenced from our mRNA library, and the proteins were identified from purified venom by mass spectrometry [5]. This provides important support for the hypothesis of HGT in *Ganaspis sp.1* venom. Additionally, we did not find significant homology to either the 18S (*Blastocystis*) or 16S (*Arsenophonus* and *Sodalis*) rRNA sequences of these species, suggesting that they are not present in *Ganaspis sp.1* wasps.

HGT into eukaryotes is a comparatively rare event, however, there are examples of horizontally transferred genes being expressed in eukaryotic transcriptomes [74], and eukaryotic HGT is most commonly observed between intracellular microbes and their hosts [75–80]. The intracellular localization of *Blastocystis spp.*, *Arsenophonus spp.* and *Sodalis spp.* therefore suggests a possible mechanism for the transfer of these genes into the *Ganaspis sp.1* genome. While the functions of these genes are unknown, their presence in *Ganaspis sp.1* venom suggests they may play a role in successful parasitization. There are multiple examples of horizontally transferred genes in arthropod genomes that are expressed and play important functional roles in metabolism and immunity [81–84]. Of particular relevance is the GH19 family of glycoside hydrolase chitinases, which is widespread in bacteria and plants but only found in one metazoan lineage, the chalcid parasitoid wasps [42]. GH19 chitinase is expressed in the venom glands of multiple chalcid parasitoid species, including the model parasitoid wasp *N. vitripennis*, where it acts to manipulate the transcription of host immune genes following infection [42].

### Paralog evolution

Along with the recruitment of *de novo* or horizontally transferred genes, venom proteins can also arise as a result of duplications of cellular genes [29–31]. The duplication of either entire or partial genes can result in the production of novel genes via paralog formation or exon shuffling [85]. We predict that these processes are likely to contribute to the evolution of the composition of *Ganaspis sp.1* venom. A venom gene formed from a genome duplication would show a high degree of similarity to an extant cellular gene. Additionally, we would predict that if the genome duplication event was specific to *Ganaspis sp.1* and/or closely related parasitoids, we would expect that the resulting genes would have only a single homolog in more distantly related species. In order to identify venom encoding genes that may have arisen as a result of recent genome duplications, we searched for venom proteins with significant homology to a cellular protein, and with only a single homolog in the *N. vitripennis* and *D. melanogaster* genomes.

We found 10 venom genes that meet these criteria and identified their paralogs among the cellular proteins (Table 1; Supplementary Table 3). Interestingly, the *D. melanogaster* homologs of 9 of these 10 venom paralogs are important for organismal viability or fertility (Table 1). Several of these *D. melanogaster* genes are involved in energy production processes including glycolysis (*Ald*), the TCA cycle (*Nc73EF*), and the electron transport chain (*Cyt*-*c*-*d* and both the alpha and beta subunits of ATP synthase, *blw* and *ATPsynB*). The remaining genes include the peptidyl-prolyl isomerase *Cyp1*, the annexin family member *AnxB9*, the aldo-keto reductase *CG10638*, the M13 family metallopeptidase *Nep2*, and the myosin light chain *Mlc2*.

**Table 1.** Predicted venom paralogous genes are shown with their fly homolog and fly mutant phenotype.

| Gene ID | Venom Paralog | Fly Homolog | Mutant Phenotype |
| --- | --- | --- | --- |
| Aldo-Keto Reductase | comp2501_c0_seq1 | *CG10638* | Viable and Fertile |
| Aldolase | comp488_c0_seq2 | *Ald* | Lethal |
| Annexin B9 | comp829_c0_seq1 | *AnxB9* | Partially Lethal |
| ATP Synthase Alpha | comp448_c0_seq1 | *blw* | Lethal/Male Sterile |
| ATP Synthase Beta | comp393_c0_seq1 | *ATPsynB* | Lethal/Female Sterile |
| Cyclophilin | comp148_c1_seq1 | *Cyp1* | Lethal |
| Cytochrome c | comp346_c0_seq1 | *Cyt-c-d* | Male Sterile |
| Myosin Light Chain | comp1347_c0_seq1 | *Mlc2* | Lethal |
| Neprilysin | comp844_c0_seq1 | *Nep2* | Partially Lethal |
| Oxoglutarate Dehydrogenase | comp1724_c0_seq2 | *Nc73EF* | Partially Lethal |

The vital roles of these genes in organismal development and fertility suggest that their functions are likely to be highly maintained by selection. If these important roles are conserved between *D. melanogaster* and *Ganaspis sp.1*, we would predict that the cellular paralogs would also be subject to purifying selection. Conversely, the venom paralog would be free to diversify through the process of neofunctionalization [85], perhaps acquiring dominant negative properties that would allow *Ganaspis sp.1* venom to manipulate these important processes in *D. melanogaster* cells. These venom proteins could thereby inhibit host immunity or alter host metabolism to confer a benefit to the developing parasitoid offspring.

In support of this hypothesis, we find that *comp1724*, the venom paralog of oxoglutarate dehydrogenase (OGDH), is lacking the dimerization interface normally found within the Transketolase PYR domain (pfam02779; Figure 3A). This dimerization interface is present in the sequences of *comp9906*, the *Ganaspis sp.1* cellular OGDH paralog, and OGDH family members in other species. It allows OGDH to interact with the other subunits of the oxoglutarate dehydrogenase complex (OGDC), and the formation of this complex is required for enzymatic function [86,87]. This suggests that venom OGDH would not be capable of forming an enzymatically active complex. However, venom OGDH does encode the necessary sequences for binding to thiamine pyrophosphate (TPP; Figure 3B), an obligate cofactor for OGDC function [88,89], and also an essential cofactor for the pyruvate dehydrogenase complex (PDC) [90,91]. Based on these sequence data, we would predict that in host cells, venom OGDH would have a dominant negative effect by sequestering available TPP into nonfunctional interactions. This would likely result in altered host metabolism through inhibiting the function of the OGDC and PDC complexes.

**Figure 3.**
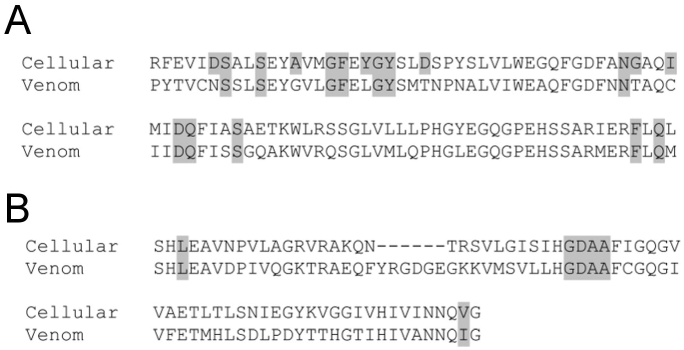
Oxoglutarate dehydrogenase sequence alignments. (A,B) Sequence alignments for the (A) Transketolase PYR domain and (B) Thiamine pyrophosphate binding domain from *comp9906*, the non-venom (Cellular) oxoglutarate dehydrogenase paralog, and *comp1724*, the venom paralog of oxoglutarate dehydrogenase (Venom). Functional residues are highlighted, demonstrating the specific lack of conservation of functional residues in the Transketolase PYR domain in the venom paralog.

### Conserved proteins and venom specific isoforms

In addition to the presence of venom specific proteins, it has previously been demonstrated that multiple venom protein encoding genes are also expressed outside the venom gland [29,40]. This suggests that they encode multifunctionalized proteins that likely act both within the venom and in non-venom cellular processes. We have found that one venom encoding gene, *Ganaspis sp.1* SERCA, encodes two protein isoforms (SERCA_1002_ and SERCA_1020_). We demonstrated that SERCA_1020_ is found throughout the wasp while SERCA_1002_ is found only in the venom [5], suggesting the presence of a venom-specific SERCA isoform. A similar phenomenon is seen in the parasitoid wasp *Pteromalus puparum*, the venom of which contains a venom-specific isoform of the SERPIN domain protein *Pp*S1V [41]. In our current analysis we identified 31 venom proteins that are encoded by genes with multiple isoforms in the *Ganaspis sp.1* transcriptome (Supplementary Table 4). We would hypothesize that the identified venom transcripts may correspond to venom-specific isoforms (as in the case of SERCA and *Pp*S1V), however, this hypothesis will require additional testing.

### Functional Annotation of *Ganaspis sp.1* Venom Proteins

Parasitoid venom proteins are capable of altering host physiology, metabolism and immunity, and while there have been numerous studies into the roles of individual venom proteins [5–7,92,93], we wanted to explore the predicted activity of the *Ganaspis sp.1* venom proteome taken as a whole. Interestingly, we find that *Ganaspis sp.1* venom genes encode a wide range of putative functions, with 863 NCBI-curated conserved domain families and superfamilies and an additional 241 PFAM domains identified from the 166 venom proteins. Notably, we find that 14 of these conserved domains are significantly enriched in the venom proteome when compared with our random subset of cellular proteins (Table 2). This suggests that the *Ganaspis sp.1* venom proteome may be composed of a distinct functional set of proteins.

**Table 2.**
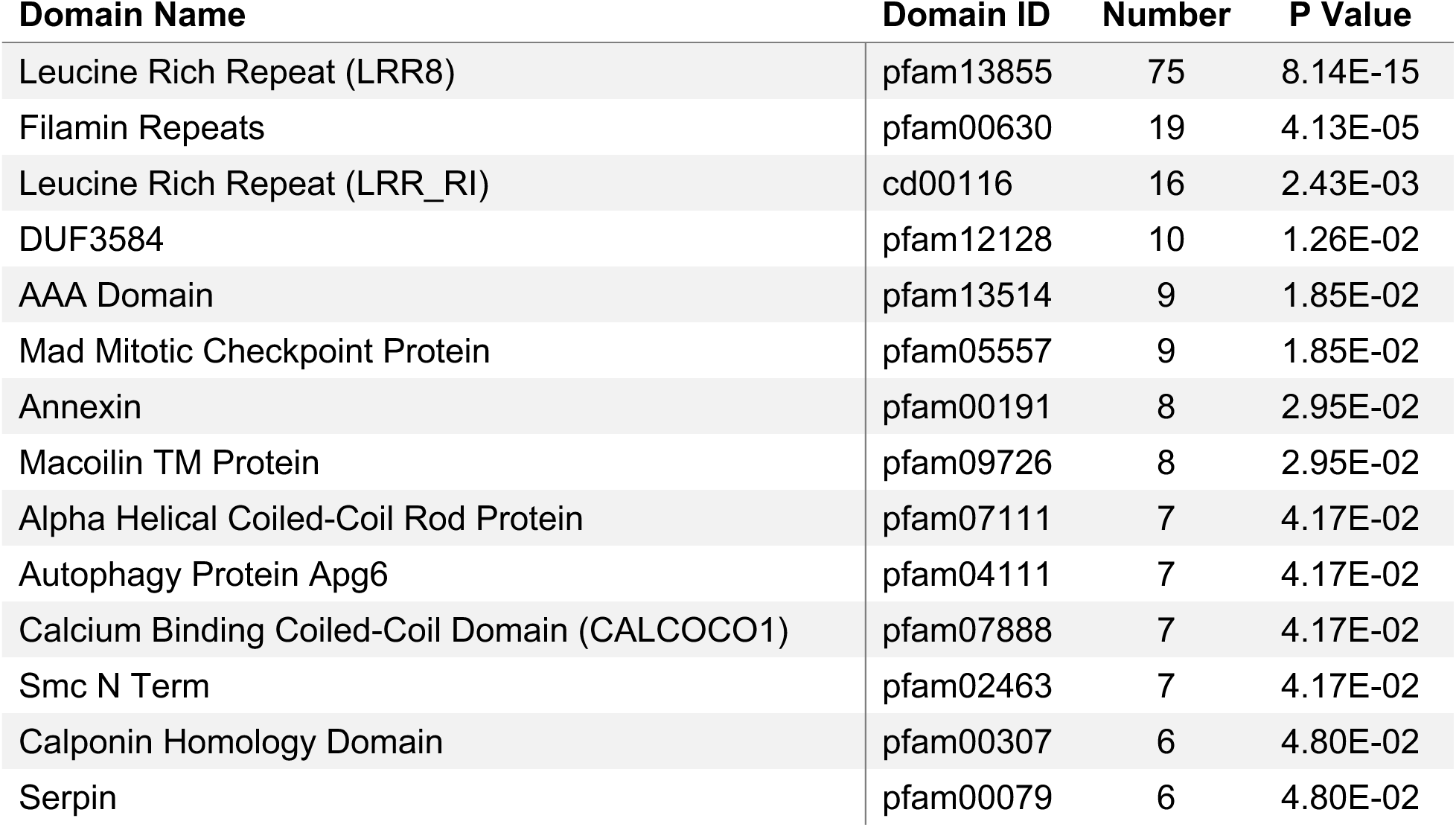
Conserved domains showing enrichment in the venom protein sample.

The list of enriched functional domains provides insight into the immunomodulatory activity of *Ganaspis sp.1* venom. We have previously shown that *Ganaspis sp.1* venom blocks the activation of host immune cells by antagonizing intracellular calcium signaling via activity of the venom specific SERCA_1002_ isoform [5]. SERCA is a calcium ATPase that transports calcium ions out of the cytosol [94], and the addition of *Ganaspis sp.1* venom significantly decreased calcium ion levels in *D. melanogaster* immune cells [5]. Our venom analysis suggests that *Ganaspis sp.1* venom may contain additional proteins with calcium regulatory potential, as revealed by the enrichment of the CALCOCO1, Calponin Homology, and Annexin calcium interacting domains [95–97]. The roles of these proteins in *Ganaspis sp.1* are unknown, but raise the possibility that multiple mechanisms act to antagonize host calcium signaling.

Additionally, insects make extensive use of serine protease cascades in triggering immune responses [98–100], and in turn, these responses are negatively regulated by serine protease inhibitor (SERPIN) domain containing proteins [101,102]. Many previously investigated parasitoid wasps have been found to contain SERPIN domain proteins in their venoms [7,29,31,41,103] suggesting that these parasitoids may use SERPIN domain proteins to negatively regulate host immunity. Interestingly, we also found multiple SERPIN domain containing proteins in *Ganaspis sp.1* venom, suggesting that this might be a shared feature of parasitoid venoms.

Along with these known venom functions, our proteome-wide analysis has revealed potential novel venom functions. The most abundant of the enriched domains are predicted to mediate protein-ligand interactions. These domains include two distinct forms of Leucine Rich Repeats (LRR; LRR8 repeats [pfam13855] and LRR_RI repeats [cd00116]) and Filamin Repeats (FR; pfam00630). LRRs are 20-29 amino acid long motifs that form discrete helical structures and allow for binding to specific partners in both homotypic and heterotypic interactions [104,105]. Similarly, FR motifs mediate a wide range of protein-protein interactions through the formation of an immunoglobulin-like beta sandwich structure [106–108]. This suggests that the *Ganaspis sp.1* venom proteins containing these domains may be able to interact with specific host proteins following infection.

### Evolutionary minimization of *Ganaspis sp.1* venom proteins

Protein minimization is a concept originally described in synthetic biology, in which proteins are artificially reduced to the minimum sequence that will support a selected function [109,110]. In our analysis, we found a subset of 17 venom proteins that appear to represent evolutionarily minimized proteins (Table 3). In comparison with the homologous proteins in *D. melanogaster*, these “minimized” venom proteins are smaller (with an average size of 37.3 kDa for the venom proteins compared to an average size 127.9 kDa for the homologs, p= 3.559e-04) and encode a decreased number of total conserved domains (3.0 vs. 10.9 total conserved domains per protein, p = 3.961e-05). We find that 14 of these minimized proteins contain conserved LRR domains and are therefore predicted to be involved in protein-ligand interactions. The other minimized proteins contain CUB (cd00041), EGF/CA (pfam07645) or LIM (cd08368) family conserved domains, all of which are also predicted to mediate protein binding [111–114].

**Table 3.** Minimized venom proteins and their *Drosophila* homologs listed by principal conserved domain, size and number of distinct and total conserved domain. *Drosophila* homologs with a known role in immunity are indicated.

| Domain | Venom | Size (kDa) | Distinct CD | Total CD | Homolog | Size (kDa) | Distinct CD | Total CD | Immune Role |
| --- | --- | --- | --- | --- | --- | --- | --- | --- | --- |
| LRR | comp3727_c1_seq2 | 69.8 | 1 | 6 | 18 wheeler | 154.8 | 3 | 13 | [121,122] |
| LRR | comp6788_c0_seq1 | 40.9 | 1 | 5 | artichoke | 172.1 | 2 | 15 |  |
| EGF | comp248_c0_seq1 | 24.8 | 1 | 1 | CG31999 | 102.1 | 2 | 12 |  |
| CUB | comp282_c0_seq1 | 15.3 | 1 | 1 | CG42255 | 403.5 | 2 | 25 |  |
| LRR | comp1786_c0_seq1 | 50.3 | 1 | 4 | CG7896 | 154.1 | 3 | 18 |  |
| LRR | comp2027_c0_seq2 | 35.5 | 1 | 1 | kek6 | 93.4 | 3 | 4 | [128,129] |
| LRR | comp2027_c0_seq22 | 9.4 | 1 | 1 | kek6 | 93.4 | 3 | 4 | [128,129] |
| LRR | comp4656_c0_seq11 | 21.2 | 1 | 2 | tartan | 81.9 | 2 | 7 |  |
| LRR | comp2027_c0_seq1 | 33.1 | 1 | 4 | tartan | 81.9 | 2 | 7 |  |
| LRR | comp13781_c0_seq1 | 90.1 | 1 | 8 | Toll | 124.6 | 3 | 11 | [123,124] |
| LRR | comp3727_c1_seq1 | 89.0 | 1 | 4 | Toll-7 | 160.9 | 3 | 14 | [126,127] |
| LRR | comp4656_c0_seq2 | 38.7 | 1 | 5 | Toll-7 | 160.9 | 3 | 14 | [126,127] |
| LRR | comp6082_c0_seq1 | 32.2 | 1 | 2 | CG1504 | 46.7 | 1 | 1 |  |
| LRR | comp2027_c0_seq10 | 22.4 | 1 | 1 | CG32055 | 61.1 | 1 | 6 | [125] |
| LRR | comp2027_c0_seq4 | 34.0 | 1 | 3 | chaoptin | 152.0 | 1 | 15 |  |
| LRR | comp2027_c0_seq6 | 18.0 | 1 | 2 | Connectin | 77.2 | 1 | 7 | [129] |
| LIM | comp934_c0_seq1 | 8.3 | 1 | 1 | Mlp84B | 53.5 | 1 | 5 |  |

The minimized *Ganaspis sp.1* venom proteins also show reduced domain complexity when compared to their *D. melanogaster* homologs; the *D. melanogaster* proteins have an average of 2.1 distinct conserved domains per protein, whereas the minimized *Ganaspis sp.1* venom proteins all contain only a single type of conserved domain (Table 3; p = 3.103e-05). More specifically, 12 of the proteins are missing domains that are present in the host homologs, including both conserved functional domains and predicted transmembrane domains (Figure 4). These alterations in protein domain architecture are often seen in dominant negative protein isoforms [115–118]. We propose that these minimized venom proteins may act as dominant negatives by directly binding to, and thereby sequestering, host proteins. This dominant negative strategy is also widely used by bacterial and viral pathogens, where pathogen proteins inhibit host immunity by interacting with host immune factors [119–122].

**Figure 4.**
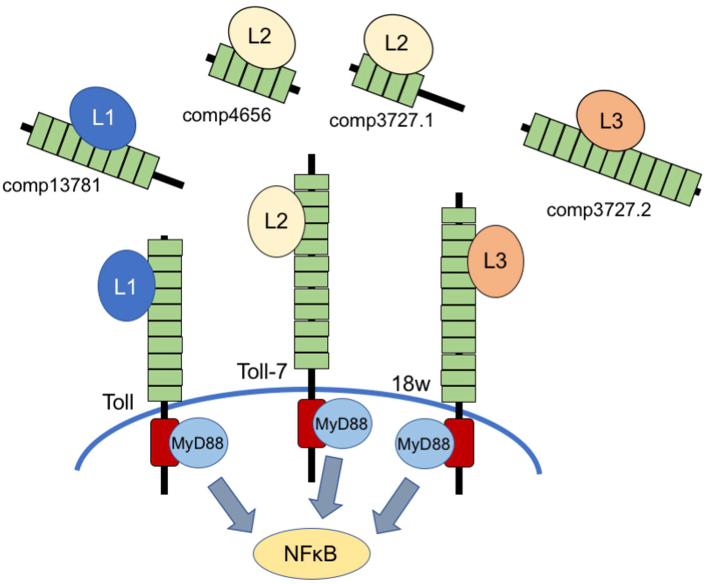
*Ganaspis sp.1* Toll-like venom proteins. A schematic diagram illustrating the differences between the *Ganaspis sp.1* Toll-like venom proteins, and their *Drosophila* homologs. The *Ganaspis sp.1* Toll-like venom proteins are smaller and have fewer LRR domains (green boxes). LRR domains will predicted homologous ligand affinities are shown binding to the same hypothetical ligands (L1, L2, L3). The Toll-like venom proteins additionally do not contain predicted transmembrane domains or TIR domains (red boxes), and so would not be predicted to signal through the MyD88 protein to potentiate NFκB activity.

Indeed, several of the *D. melanogaster* homologs are involved in immune responses (Table 3, [123–131]) and we would therefore predict that interfering with activity of these genes would aid *Ganaspis sp.1* in evading host immunity. Three of these host immune genes encode Toll-like receptors (TLRs: *Toll* [*Tl*], *Toll*-*7* and *18 wheeler* [*18w*]). *D. melanogaster* TLRs are transmembrane proteins, and receive signals by the binding of ligands to their extracellular LRR domains [132,133]. The signal is then transduced through the interaction of the intracellular Toll/Interleukin-1 Receptor homology domain (TIR) with the downstream adaptor protein MyD88, leading to the activation of NFκB transcription factors and subsequent immune signaling [134,135]. The venom homologs of these TLRs have highly conserved LRRs, but lack transmembrane or TIR domains (Figure 4). We would predict that these proteins would bind to endogenous TLR ligands, but would be unable to activate downstream signaling and could act to sequester ligands and subsequently dampen host signaling in response to infection.

## Conclusions

Our bioinformatic analysis reveals the complexity of the evolution and function of parasitoid venoms. First, our results suggest that the evolution of venom composition in a single parasitoid species is driven by multiple mechanisms. *Ganaspis sp. 1* venom encoding genes show evidence of gene multifunctionality and co-option, along with neofunctionalization following gene duplication. Additionally, several of these venom genes appear to have evolved as *de novo* or horizontally transferred genes. Second, our analyses suggest several potential virulence functions of *Ganaspis sp.1* venoms. The enriched functional domains found in these proteins suggest that venoms may act to inhibit fly immunity through antagonizing calcium signaling dependent immune cell activation or blocking serine protease cascades. Furthermore, we have uncovered evidence suggesting that the venom paralogs arising from recent gene duplication events may have evolved dominant negative traits that would interfere with essential host functions. Finally, we have identified a subset of venom proteins that appear to have undergone a minimization process resulting in smaller and less complex venom proteins that we would predict could interfere with host immune function by sequestering immune receptor ligands. Testing the predictions we have made based on our analyses will provide further insight into the function of parasitoid venoms, and open new areas of research within this exciting field.

## Acknowledgements

We would like to thank Drs. Jeff Helms and Pennapa Manitchotpisit for their assistance. This research did not receive any specific grant from funding agencies in the public, commercial, or not-for-profit sectors, and was supported by the School of Biological Sciences at Illinois State University.

